# BEHAVIOURAL AND MOLECULAR EFFECTS OF CANNABIDIOLIC ACID IN MICE

**DOI:** 10.1101/2020.06.22.165076

**Authors:** Laia Alegre-Zurano, Ana Martín-Sánchez, Olga Valverde

## Abstract

**Aims:** Cannabidiolic acid (CBDA) is one of the most abundant phytocannabinoid acids in the *Cannabis Sativa* plant. It has been shown it is able to exert some therapeutic effects such as antiemetic, anti-inflammatory, anxiolytic or antidepressant, although some of them remain under debate. In the present study we aim to assess the potential effects of CBDA on different behaviours and on the modulation of neuroinflammatory markers in the prefrontal cortex (PFC).

**Main methods:** the effects of acute and/or chronic CBDA (0.001-1 mg/kg i.p.) treatment were evaluated on cognitive, emotional, motivational and nociceptive behaviours in male CD1 mice. For this, Y-maze and elevated plus maze paradigms, spontaneous locomotor activity, social interaction, hot-plate, formalin and tail suspension tests were used. We also studied the effects of CBDA on the rewarding responses of cocaine in the conditioned place preference (CPP) paradigm. PFC was dissected after acute and chronic CBDA treatments to evaluate inflammatory markers.

**Key findings:** acute CBDA treatment induced antinociceptive responses in the hot-plate test. A 10-day chronic CBDA treatment reduced despair-like behaviour in the tail suspension test. CBDA did not alter the remaining behavioural tests assayed, including cocaine-induced reward in the CPP. Regarding the biochemical analysis, chronic CBDA treatment diminished peroxisome proliferator-activated receptor gamma (PPAR-γ) and increased interleukin-6 (IL-6) protein levels in PFC.

**Significance:** these results show that CBDA has limited *in vivo* effects modulating mice behaviour, highlighting the current disagreement regarding its therapeutic potential.

## INTRODUCTION

The *Cannabis Sativa* plant produces more than 80 different phytocannabinoids. Interestingly, the natural form of most of them is the acid form [1]. The first cannabinoid acid discovered was the cannabidiolic acid (CBDA) [1], which is one of the most abundant phytocannabinoid acids [2]. During decades, the study of phytocannabinoids has largely ignored their acid forms and so, knowledge about such is very limited. However, recently there has been a rise in the number of studies focusing on CBDA’s potential targets and pharmacological effects [3].

*In vitro* research of the mechanisms of action of CBDA points it out as a possible selective inhibitor of the proinflammatory enzyme ciclooxigenase-2 (COX-2) [4,5] although other studies failed in replicating these results [6]. Additionally, CBDA could act as an enhancer of the receptor of serotonin 1A (5-HT_1A_) activation [7–9], an agonist of the transient receptor potential cation channel vanilloid 1 (TRPV1) and transient receptor potential cation channel ankyrin 1 (TRPA1), and an antagonist of transient receptor potential cation channel melastatin 8 (TRPM8) revealed by *in vitro* assays [10,11]. Recently, CBDA has been found to act as a dual proliferator-activated receptor (PPAR) α/γ agonist in computational [12] and *in vitro* [12,13] cell culture experimental approaches, including mouse and human brain cells.

So far, very few studies have addressed the *in vivo* pharmacological responses of CBDA. Bolognini et al. (2013) found that CBDA (0.1 and 0.5 mg/kg) displays antiemetic effects for nausea from different aetiology, probably due to the enhancement of 5-HT_1A_ activation [7]. Such effect have been replicated in other studies [8,9,14–16]. Regarding the putative CBDA effects on anxiety, some authors reported that CBDA induced anxiolytic-like effects in rats in the open-field test using CBDA (0.5 and 5 mg/kg) [15], whereas others could only replicate such effect in previously stressed rats in the light-dark emergency test with doses between 0.1 and 100 mg/kg, but not in basal conditions [17]. Recently, Hen-Shoval et al. (2018) found that an acute oral administration of CBDA methyl ester (HU-580) reduces despair-like behaviour in forced swimming test, using Wistar-Kyoto (WKY) and Flinders Sensitive Line (FSL) rat, which represent genetic models of depression [18]. A recent study showed that CBDA (0.01 mg/kg) reduces the carrageenan-induced hyperalgesia and oedema in a rodent model of inflammatory pain, by a TRPV1-mediated mechanism [19]. Lately, Anderson et al. (2019) found anticonvulsant effects for CBDA at a dose of 10 mg/kg in a mouse model of Dravet Syndrome [20]. Taken together, these studies suggest CBDA as a promising therapeutic strategy while evincing the lack of enough scientific insights regarding its pharmacological effects.

During the last decades, phytocannabinoids are gaining more attention in the field of drug addiction due to their modulatory role on other neurotransmitter and modulatory systems involved in the control of reward and motivation [21,22]. Nevertheless, to our best knowledge, there are no studies in the literature focusing on the possible modulatory effects of phytocannabinoid acids in the rewarding effects of psychostimulants.

Therefore, the main aims in the present investigation were: (i) to study the behavioural effects of CBDA after acute and chronic treatments in male mice; (ii) to assess the levels of PPAR-γ and interleukin-6 (IL-6), inflammatory markers, after acute and chronic treatments of CBDA; and (iii) to examine the effects of CBDA on the rewarding effects of cocaine evaluated in the CPP.

## MATERIALS AND METHODS

### Animals

Male CD1 mice (postnatal day 35) were purchased from Charles River (Barcelona Spain) and transported to our animal facility (UBIOMEX, PRBB). Animals were maintained in a 12-hour light-dark cycle. Mice were housed at a stable temperature (22°C±2) and humidity (55% ± 10%), with food and water *ad libitum*, and they were allowed to acclimatize to the new environmental conditions for at least five days prior to the experiments. All animal care and experimental protocols were approved by the UPF/PRBB Animal Ethics Committee (CEEA-PRBB-UPF), in accordance with European Community Council guidelines (2016/63/EU).

### Drugs

CBDA (0.001; 0.01; 0.1; 1 mg/kg, i.p.) was kindly provided by Phytoplant Research S.L., Córdoba, Spain. For the preparation of the solution, CBDA was dissolved in ethanol, cremophor and 0.9% NaCl (1:1:18). Cocaine HCl (10 mg/kg i.p.) was purchased in Alcaliber S.A., Madrid, Spain, and was dissolved in 0.9% NaCl. The volume of injection for both drugs was 0.1 mL per 10 g of mouse body weight.

### Drug administration protocols and groups

CBDA was administered acutely and using a 10-day chronic treatment. The experimental groups (9-10 mice/group) were VEH, CBDA 0.001 mg/kg, CBDA 0.01 mg/kg, CBDA 0.1 mg/kg and CBDA 1 mg/kg (acute or chronically administered) for each experiment. In order to reduce the number of animals used, some sets of animals underwent two different experiments when behavioural test specificities allowed it.

The acute treatment consisted on the administration of CBDA 30 minutes before starting the behavioural test (Fig 1a), then, specific behaviours were evaluated in the different behavioural procedures. For the chronic treatment, CBDA was administered during 10 consecutive days and 24 hours after the last CBDA administration, mice were evaluated in the elevated plus maze (EPM) and the Y-maze (Fig 1b). In addition, another set of animals underwent a stress exposure session, followed by the elevated plus maze and the tail-suspension test (TST), as showed in Figure 1c.

**SINGLE COLUMN Fig 1.**
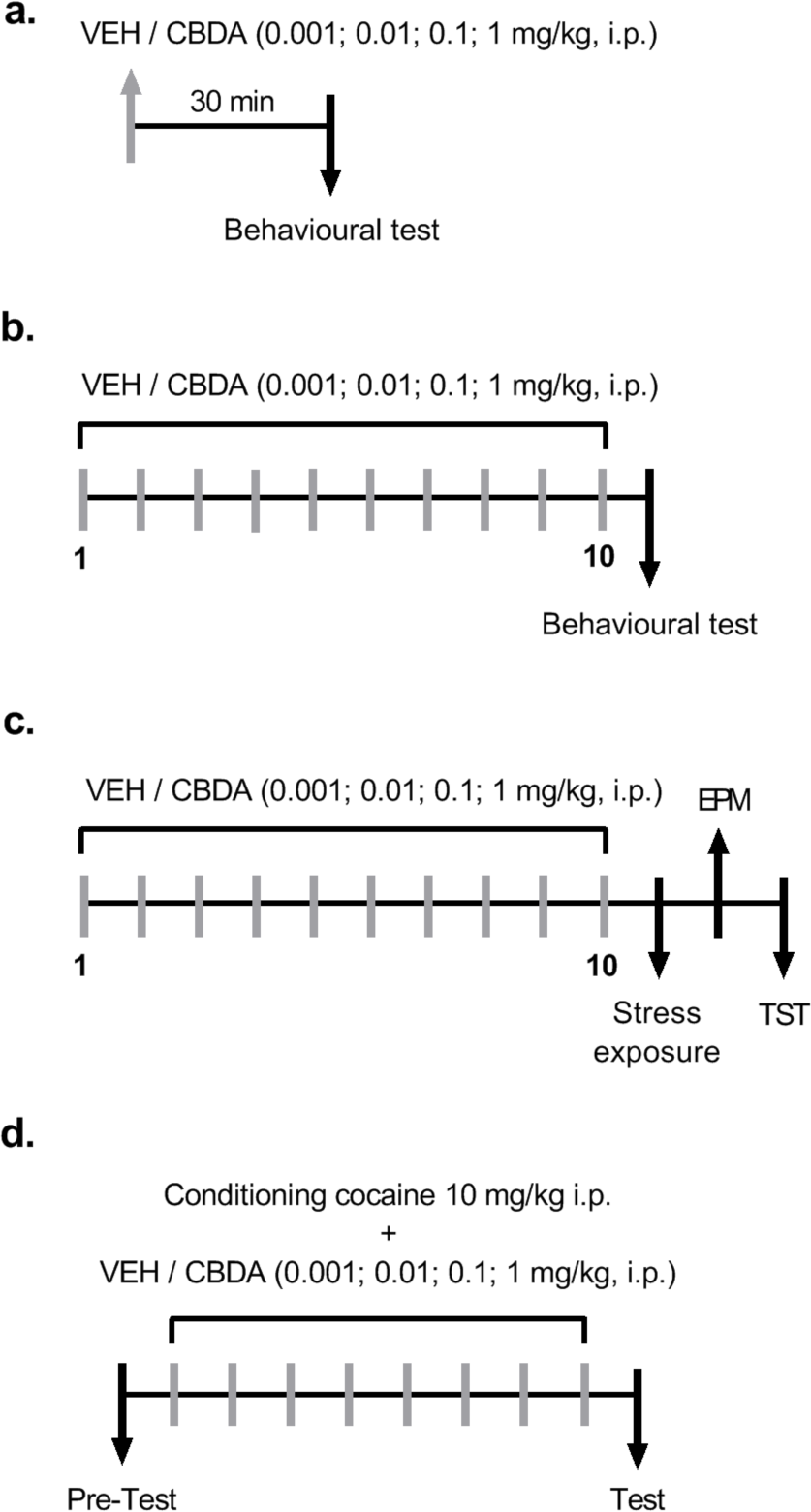
Schematic description of the different protocols used. (a) Acute CBDA treatment was administered 30 minutes before the behavioural test and (b) chronic treatment was administered daily for 10 days. Behavioural test started 24 hours after the last CBDA injection. (c) 24 hours after a 10-days chronic treatment of CBDA, mice underwent stress session, and TST 48 hours later. (d) During cocaine-induced conditioned place preference, CBDA treatment was administered 30 minutes before every session during the eight days of conditioning.

A different set of animals was used for cocaine-induced conditioned place preference (CPP). The groups were VEH-Saline, VEH-Cocaine 10 mg/kg, CBDA 0.01 mg/kg-Saline, CBDA 0.01 mg/kg-Cocaine 10 mg/kg, CBDA 0.1 mg/kg-Saline, and CBDA 0.1 mg/kg-Cocaine 10 mg/kg. In this case, CBDA was administered daily during the 8-days conditioning phase 30 minutes before entering the box (Fig 1d).

### Spontaneous Locomotor Activity

Mice were evaluated in locomotor activity boxes (24 × 24 × 24 cm) (LE8811 IR, Panlab s.l.u., Barcelona, Spain) provided with 14 axes (Y and X) in a low-luminosity room. Horizontal (deambulations) and vertical (rearing) movements were automatically recorded for 45 min.

### Social interaction test

The test was conducted as previously reported [23]. Briefly, the test consisted of three 10-minute phases: habituation, sociability and social novelty. Time spent in each chamber for each phase was recorded by the SMART software (Panlab s.l.u., Spain) for subsequent analysis.

### Hot-plate test

Nociceptive responses were measured with the hot-plate test which was conducted as previously reported [24]. Two different nociceptive thresholds were evaluated: the latency (s) to lick the hind paws and to jump.

### Formalin test

The procedure was adapted from Simonin et al. (1998). The time licking or biting the hind paw after a 20 µl of 5% formalin injection was recorded during the early (minutes 0 to 5) and late phase (minutes 15 to 30).

### Y-Maze test

Mice were assessed for spatial working memory as previously reported [25]. Briefly, the maze consisted of a Y-shaped maze with three arms. The score was obtained by dividing the total number of alternations (three consecutive choices of three different arms) by the total number of choices minus 2.

### Elevate plus maze

The EPM was performed to evaluate anxiety-like behaviour in mice [24,26]. Mice were placed on the centre of the maze for 5 minutes and the number of entries and time spent in the open and closed arms were recorded by automated tracking software and analysed as follows: percentage of time spent in the open arms = (time spent in the open arms / summation of time spent in the open and closed arms) x 100.

### Tail-suspension test

Mice were individually suspended from the tip of the tail, 50 cm above a bench top for a 6-minute period. Immobility time was measured as previously reported [23].

### Stress exposure

A group of animals that received a 10-days chronic treatment of CBDA (0.001; 0.01; 0.1; 1 mg/kg) or vehicle (i.p.) were subjected to a stressful session 24 hours after the last CBDA or vehicle administration. In this session, animals were habituated for 5 minutes to the mouse operant chambers (Model ENV-307A-CT, Medical Associates, Georgia, VT, USA). Afterwards, they underwent a 10-minute session with 20 0.5 mA foot shocks of one second long at variable interval. 24 and 48 hours later, mice underwent the EPM and the TST, respectively.

### Cocaine-induced conditioned place preference

The test was performed as previously reported [26]. Briefly, during the pre-test and test, mice were placed in the central compartment of the apparatus (Cibertec S.A., Madrid, Spain) and were left free to move through the three compartments for 20 minutes. During the conditioning phase (4 cocaine pairings, 8 days), mice received a cocaine injection of 10 mg/kg, i.p. immediately before being placed into one of the two conditioning compartments for 30 min. On the alternate days, mice were treated with a saline injection and placed in the other compartment. CBDA/vehicle was administered during the 8 conditioning days 30 minutes before the cocaine/saline injection.

### Western Blot analysis

Two different sets of animals were used for tissue extraction. Mice were euthanized by cervical dislocation. For the acute treatment set, prefrontal cortex (PFC) of 30 mice (n=6/group) were extracted 90 minutes after CBDA/vehicle injection, after the spontaneous locomotor activity test (Fig 1a). For the chronic treatment set, PFC of 30 mice (n=6/group) was extracted 24 hours after the last CBDA or vehicle injection. Samples were stored at −80°C to be processed.

The tissue was processed as previously reported [26] and proteins were loaded onto 10% polyacrylamide gels and transferred to PVDF sheets (Immobilion-P, MERCK, Burlington, USA). Membranes were blocked with BSA 5% and immunoblotted using primary antibodies for β-Tubulin, PPARγ and IL-6 overnight at 4°C. After washing with TBS-T, membranes were incubated with their respective secondary fluorescent antibodies. Protein expression was quantified using a Li-Cor Odyssey scanner (Li-Cor, Lincoln, USA).

**Table 1.**
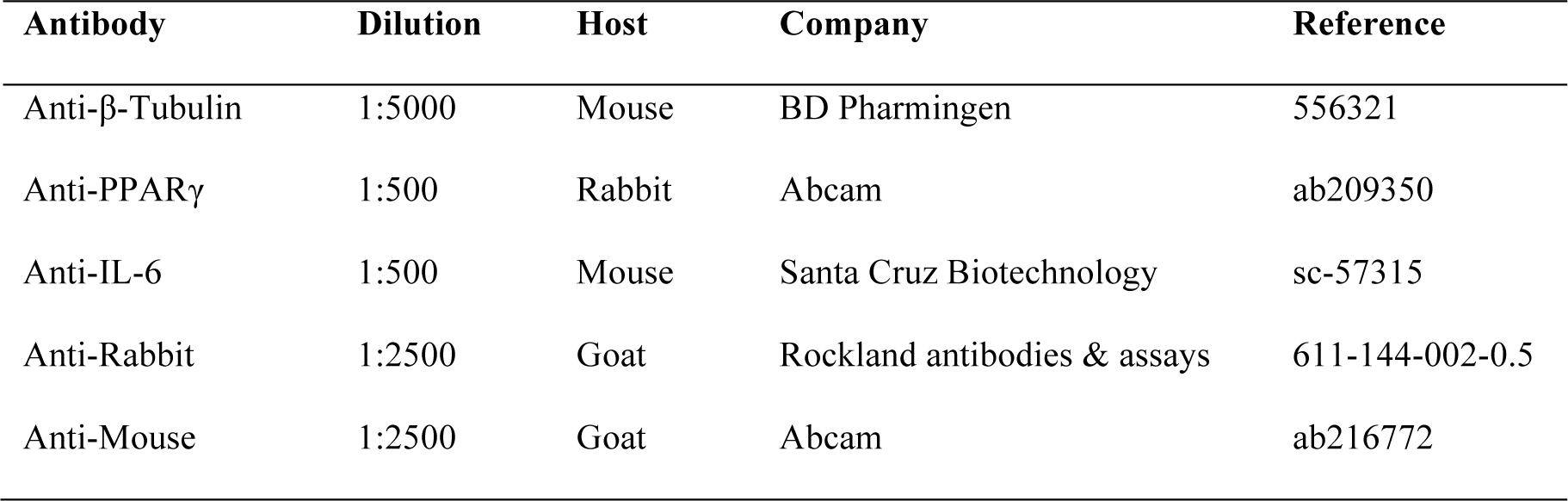
Primary and secondary antibodies used for western blot analysis

| Antibody | Dilution | Host | Company | Reference |
| --- | --- | --- | --- | --- |
| Anti-β-Tubulin | 1:5000 | Mouse | BD Pharmingen | 556321 |
| Anti-PPARγ | 1:500 | Rabbit | Abcam | ab209350 |
| Anti-IL-6 | 1:500 | Mouse | Santa Cruz Biotechnology | sc-57315 |
| Anti-Rabbit | 1:2500 | Goat | Rockland antibodies & assays | 611-144-002-0.5 |
| Anti-Mouse | 1:2500 | Goat | Abcam | ab216772 |

### Statistical analysis

Results are expressed as the mean ± SEM. Social interaction test, hot plate test, formalin test, Y-maze, EPM, TST and western blot data were analysed using a one-way analysis of variance (ANOVA) with the *CBDA treatment* as main factor. A two-way analysis of variance (ANOVA) was calculated for spontaneous locomotor activity assessment with *CBDA treatment* as a between-subject factor and *time* as a within-subject factor and for cocaine-induced CPP with *CBDA treatment* and *conditioning* as between-subject factors. When applicable, pairwise comparisons were analysed with Bonferroni’s correction. Statistical differences were found when p<0.05.

## RESULTS

### Acute CBDA treatment does not affect spontaneous locomotor activity or social behaviour but slightly modulate nociceptive responses

Regarding spontaneous locomotor activity, two-way ANOVA showed a significant effect for the *time* factor within the session for both deambulations (Fig 2a; F_8,360_=144.2; p < 0.0001) and rearings (Fig 2b; F_8,360_=19.1; p < 0.0001), without effect for *CBDA treatment* or interaction.

**1.5 COLUMN Fig 2.**
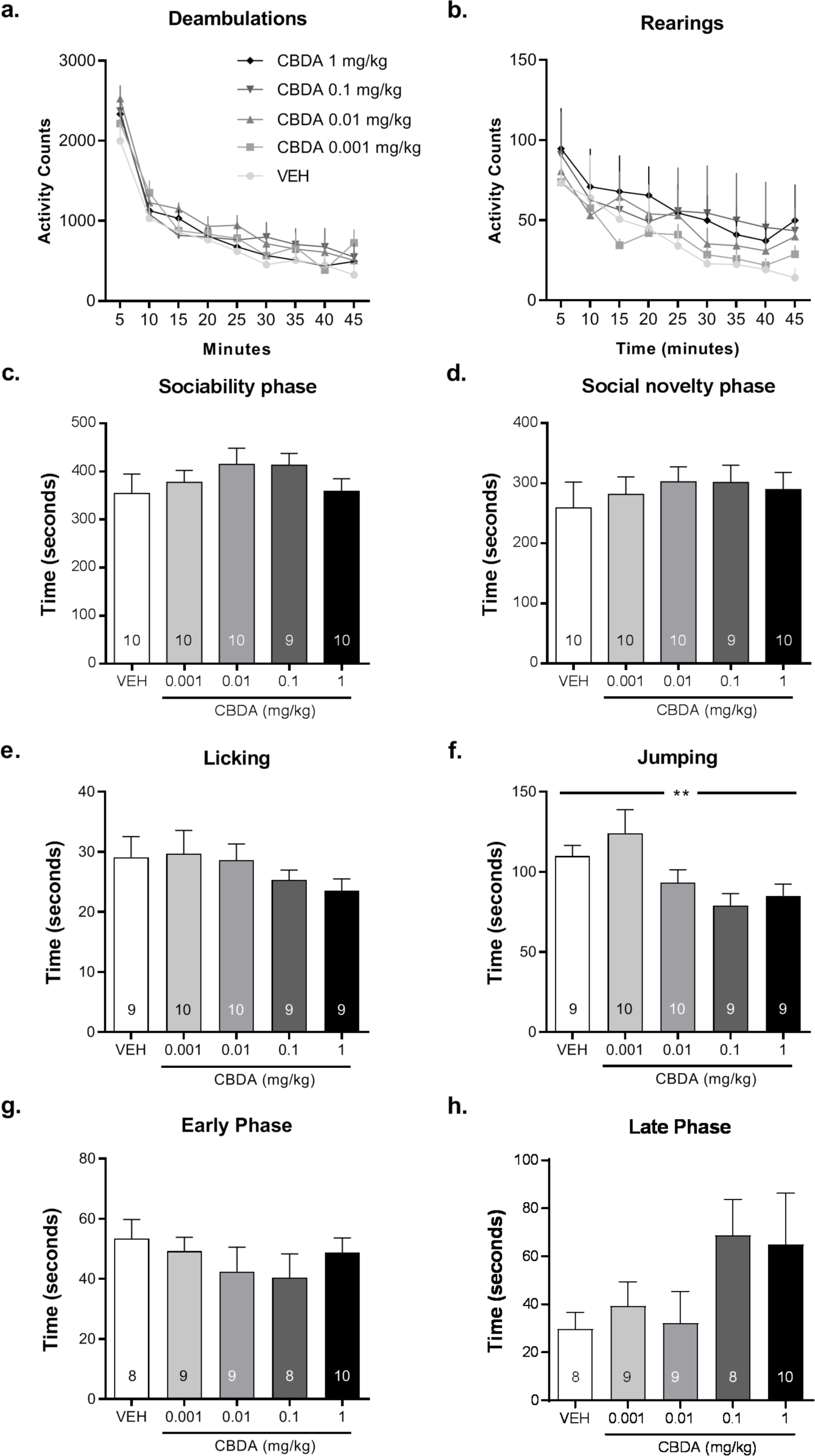
Acute CBDA treatment does not modify spontaneous locomotor activity and social behaviour, and slightly decreases supraspinal nociceptive threshold. (a) Deambulations and (b) rearings during 45 minutes after an acute VEH/CBDA treatment (Vehicle, n=10; CBDA 0.001 mg/kg, n=10; CBDA 0.01 mg/kg, n=10; CBDA 0.1 mg/kg, n=10; CBDA 1 mg/kg, n=10). Time spent in the (c) intruder / (d) new intruder compartment after acute VEH/CBDA treatment. Latency to (e) lick the paw or (f) to jump in the Hot Plate Test. Time spent licking or biting the infected paw during the (g) early or (h) late phase of the Formalin Test. One-way ANOVA, ** p < 0.01.

Considering social behaviour, one-way ANOVA analysis showed no significant effect for *CBDA treatment* in the sociability phase (Fig 2c) or in the social novelty phase (Fig 2d).

One-way ANOVA for the antinociceptive responses of CBDA in the hot-plate test revealed a significant effect in jumping behaviour (Fig 2f, F_4,42_=3.831; p < 0.01). Bonferroni’s *post hoc* comparisons showed a significant difference between CBDA 0.001 mg/kg and CBDA 0.1 mg/kg groups (p < 0.05), indicating that CBDA produced antinociceptive effects at these doses. No significant differences were found for licking behaviour for none of the assayed doses (Fig 2e). One-way ANOVA analysis for the antinociceptive responses in the formalin test, showed no effects for *CBDA treatment* in neither phase of the test (Fig 2g and Fig 2f).

### Acute or chronic CBDA treatment does not affect working memory in the Y-maze or anxiety-like behaviour in the EPM

For the Y-maze task, one-way ANOVA showed no significant differences for acute (Fig 3a) or chronic (Fig 3b) CBDA treatment.

**1.5 COLUMN Fig 3.**
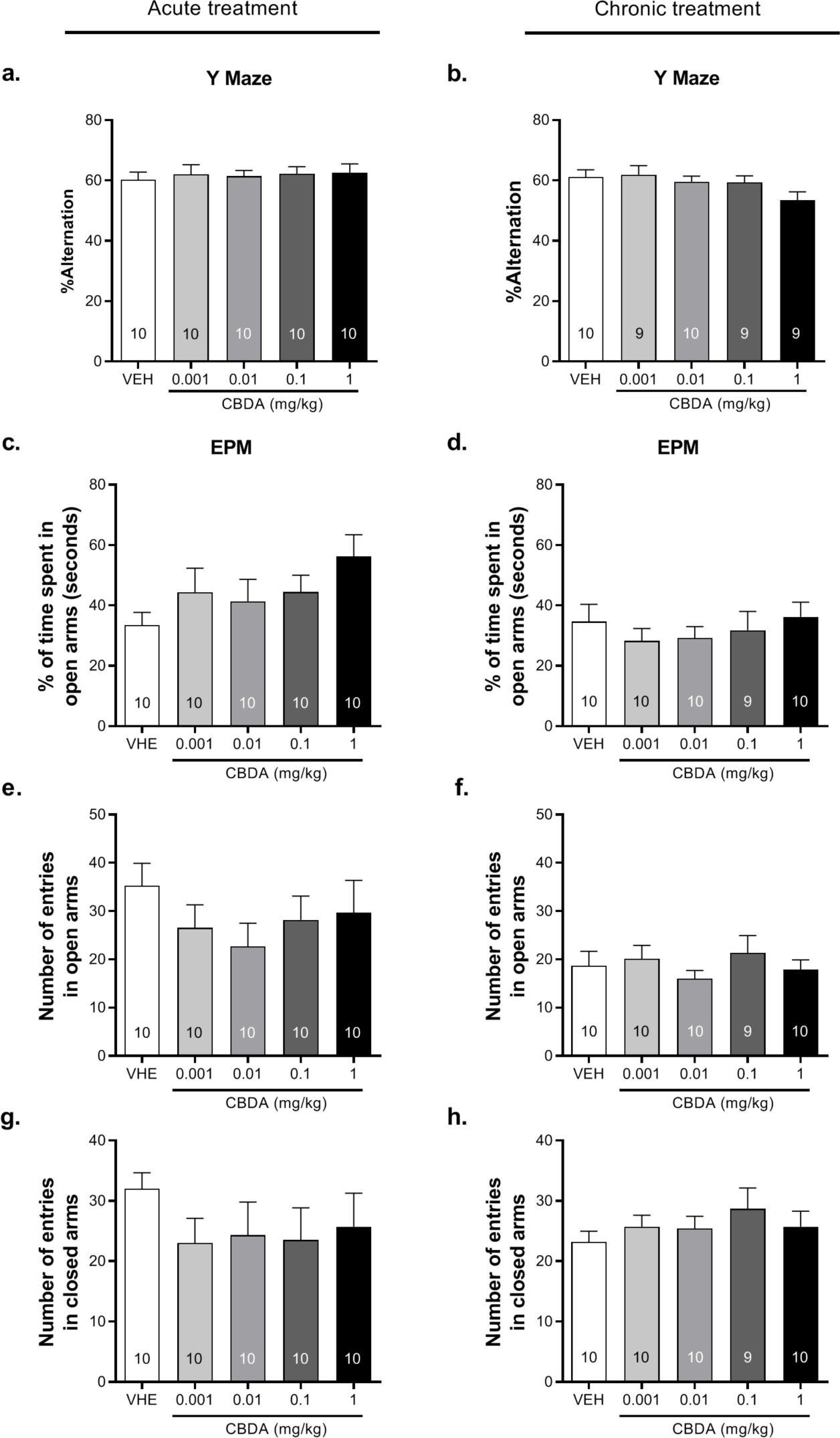
Acute or chronic CBDA treatment has no effect on working memory and anxiety-like behaviour. Percentage of spontaneous alternation in the Y-maze test after the (a) acute or (b) chronic treatment of CBDA. Percentage of time spent in open arms (c, d) and number of entries in open (e, f) and closed (g, h) arms in the EPM after an acute or chronic treatment of CBDA.

In regard to the EPM, one-way ANOVA revealed no significant differences in percentage of time spent in open arms after an acute (Fig 3c) or chronic (Fig 3d) treatment. In the same line, no differences in the number of entries in open (Fig 3e and 3f) or closed arms (Fig 3g and 3h) were found.

### Chronic CBDA treatment decreases despair-like behaviour in stressed mice

For the TST, one-way ANOVA revealed an effect of chronic *CBDA treatment* on immobility time (Fig 4a; F_4,41_=3.94; p < 0.01). Bonferroni’s *post hoc* comparisons revealed significant differences between CBDA 0.001 mg/kg and CBDA 0.01 mg/kg groups (p < 0.05) and CBDA 0.001 mg/kg and CBDA 0.1 mg/kg groups (p < 0.05) (Fig 4a), indicating that CBDA reduced despair-like effects in this behavioural model. Statistical analysis for the EPM after the stress session reported no significant effects (data not shown).

**SINGLE COLUMN Fig 4.**
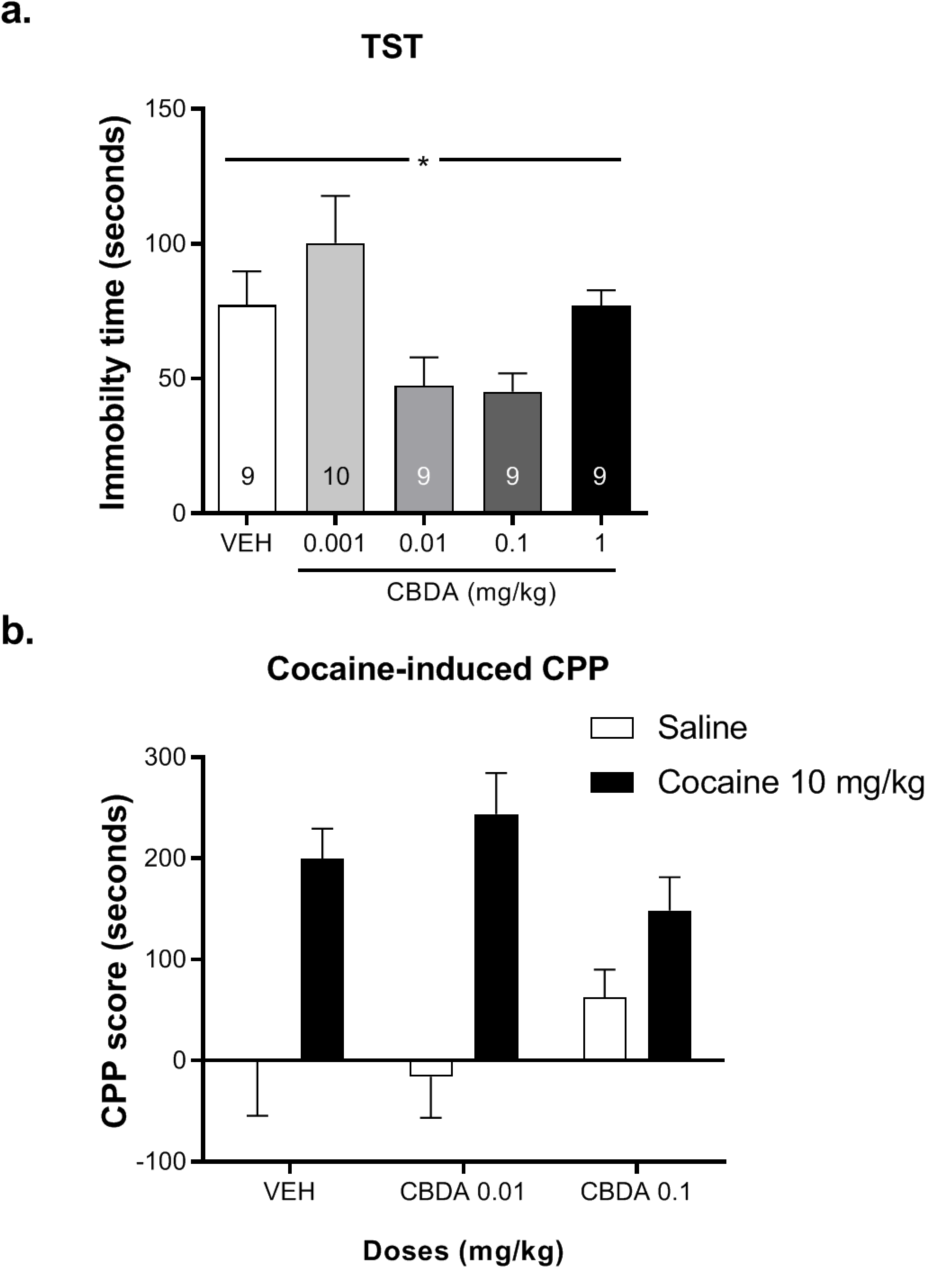
CBDA modulates despair-like behaviour but has no effect on cocaine preference. (a) Chronic CBDA treatment decreases despair-like in stressed mice. Immobility time in the TST 48 h after foot-shocks exposure. One-way ANOVA, *p < 0.05. (b) CBDA treatment administered during the conditioning pairings does not impair the preference for cocaine. The difference of time spent in the saline/cocaine-paired compartment between the test day and the pre-test day (VEH-Saline, n=9; VEH-Cocaine, n=8; CBDA 0.01 mg/kg-Saline, n=9; CBDA 0.01 mg/kg-Cocaine, n=10; CBDA 0.1 mg/kg-Saline, n=9; CBDA 0.1 mg/kg-Cocaine, n=9).

### CBDA treatment does not modify the rewarding effects of cocaine in the CPP

Two-way ANOVA showed a significant effect of cocaine in the CPP (Fig 4b; F_4,47_=31.64; p < 0.0001) for cocaine-treated groups. However, no significant effect was found for *CBDA treatment* or interaction between factors (Fig 4b).

### CBDA modifies PPARγ and IL-6 protein levels after a chronic, but not acute, treatment

A 10-day CBDA treatment (Fig 5b and 5d) showed a significant modulation of PPARγ (F_4,29_=3.215; p < 0.05) and IL-6 (F_4,31_=4.669; p < 0.01). For IL-6 protein levels, Bonferroni’s *post hoc* comparisons revealed significant differences between vehicle and CBDA 0.01 mg/kg groups (p < 0.01). However, acute CBDA treatment (Fig 5a and 5c) revealed no effect on protein levels.

**1.5 COLUMN Figure 5.**
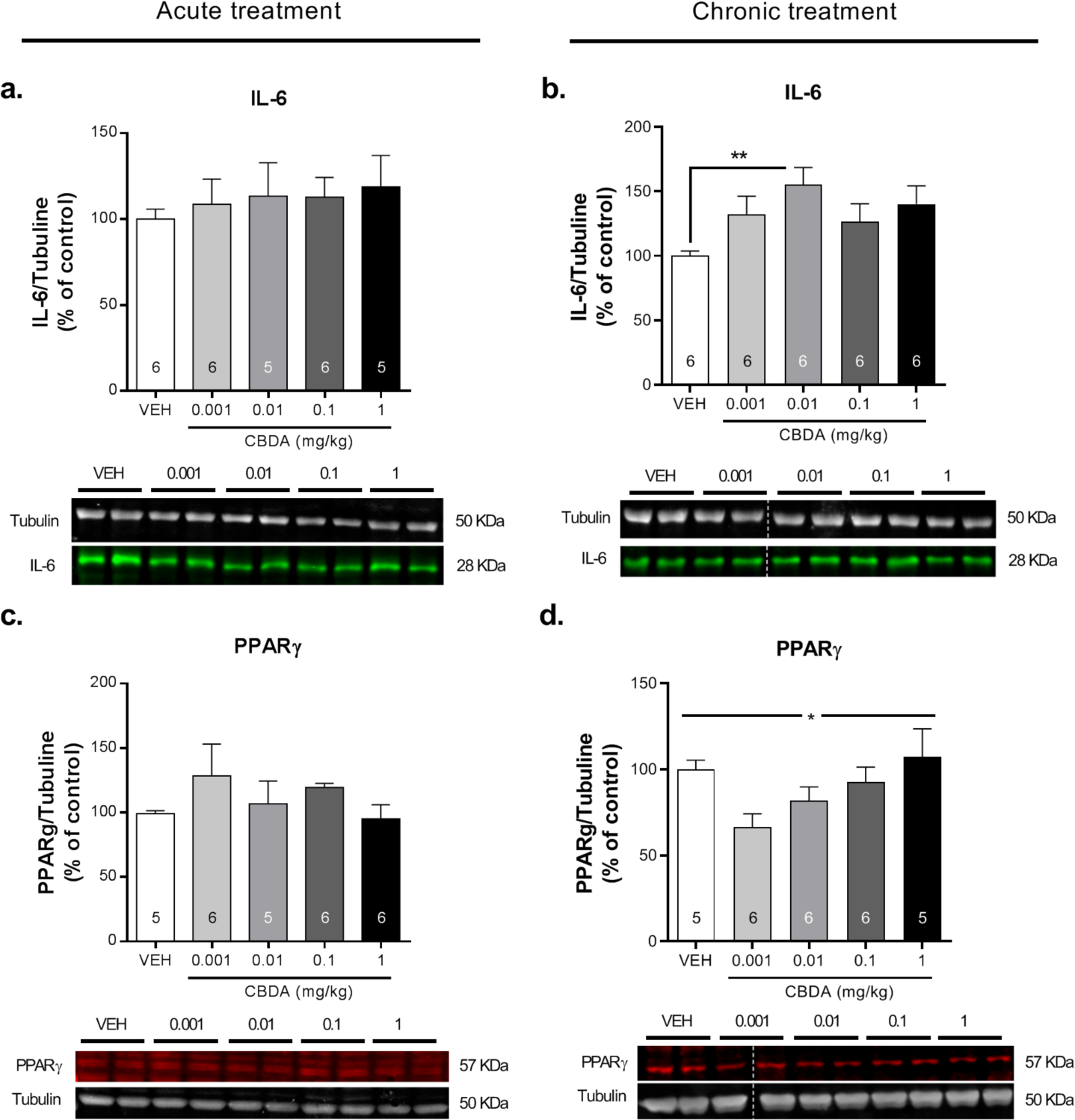
CBDA modifies PPARγ and IL-6 protein levels after a chronic, but not acute, treatment. Western blot analysis of PPARγ and IL-6 after an (a, c) acute or (b, d) chronic VEH/CBDA treatment. Bonferroni, **p < 0.01. One-way ANOVA, *p < 0.05.

## DISCUSION

Our results show that acute or chronic CBDA (0.001-1 mg/kg, i.p.) administration exerts limited effects upon mice behaviour in a wide range of behavioural tests. Acute treatment with CBDA revealed antinociceptive effects on jump latency in the hot-plate test, showing a mild modulation of the supraspinal nociceptive threshold. A 10-day chronic CBDA treatment also showed a significant attenuation on despair-like behaviour in mice exposed to the TST. Regarding the biochemical analysis, chronic CBDA treatment diminished PPAR-γ and increased IL-6 protein expression in PFC.

Consistent with our results, previous studies [14,15] showed that CBDA did not modify locomotor activity at any doses after a single administration. Therefore, we could discard that the results obtained in the behavioural tests were due to the effects of CBDA on locomotion.

CBDA shortened jump latency in the hot-plate test, revealing a modulatory role of CBDA on thermic nociceptive supraspinal responses. Nevertheless, CBDA did not show antinociceptive effects on inflammatory pain using the formalin test, suggesting a predominant role of CBDA on thermic nociceptive stimuli. Rock et al. (2018) found that CBDA 0.01 mg/kg diminished oedema and hyperalgesia of the hind paw in a rat model using the carrageenan as inflammatory agent [19]. Because the same doses were used, divergences in the outcomes might be due to differences in species, inflammatory agents and/or other parameters related to the experimental schedule. These inconsistent results highlight the need for further studies clarifying the possible role of CBDA on nociceptive control.

To our best knowledge, few studies have previously shown the effects of CBDA on anxiety-like behaviour and their results are not conclusive. In the present study, we obtained that neither acute nor chronic CBDA treatment had any effect on the anxiety-like behaviour displayed in the EPM. In agreement, acute treatments with CBDA have been found to lack anxiolytic effects in the open-field [15], the light-dark box [15,17] and conditioned freezing to a shock-paired tone [14]. Interestingly, Brierley et al. (2016) described that anxiogenic-like effects induced by the 5-HT_1A_ receptor antagonist WAY-100,635 in the novelty-suppressed feeding test were completely abolished by acute CBDA (5 mg/kg) [15]. In this last study, CBDA alone did not produce any anxiety effect, in agreement with our data, and solely reversed a previously induced anxiogenic state. In addition, also consistent with our results, Rock et al. (2017) showed no effects on anxiety-like responses in the light-dark box after a 20-days chronic CBDA treatment. However, acute CBDA (0.1-100 mg/kg) reversed anxiogenic effects of electric foot shocks in rats [17]. In our study, mice were treated with CBDA during 10 days and stressed by applying electric foot shocks to further undergo an EPM and a TST; 24 and 48 hours later, respectively. However, CBDA did not modify anxiety-related behaviours in the EPM after stress exposure. In agreement, Pertwee et al. (2018) revealed that CBDA did not show anxiolytic effects in the light-dark box after a foot shock session [9]. These findings together with our results would suggest that CBDA exerts very limited, modulatory effects upon anxiety-like behaviour.

Regarding responses in the TST, CBDA reduced despair-like behaviour 48 hours after the stress session. To our knowledge, this is the first study demonstrating such an effect of CBDA. Only a recent study published by Hen-Shoval et al. (2018) found a similar result using the forced swimming test and an acute treatment of the cannabidiolic acid methyl ester (HU-580) [18].

In regard to cocaine-induced CPP, CBDA (00.1-0.1 mg/kg) did not alter the rewarding effects of cocaine in this paradigm. This outcome remarks the differences found between CBDA and its decarboxylated form, CBD, as the last one has already been proved to decrease cocaine’s rewarding effects in the same paradigm [26]. Remarkably, CBD is currently administered at doses around 100 times higher than CBDA [7,8].

In the biochemical analysis, we assayed CBDA as an anti-inflammatory candidate as it has been found to exert such effects *in vitro* [4,5] and *in vivo* [19]. Furthermore, CBDA exerts agonistic effects on TRPV1 [10,11] and PPARγ [12,13], which are tightly related to inflammatory processes. Therefore, we measured IL-6 and PPARγ protein levels in PFC, a brain area constitutively engaged in evaluated phenomena, such as cognition, emotion, nociception and motivation [27]. Surprisingly, Western Blot analysis revealed an increase in IL-6 protein levels in the PFC after a chronic CBDA treatment, reaching significance at the dose of 0.01 mg/kg, while PPARγ showed a decreasing trend also at the lower doses tested. These results contrast with previous findings proposing CBDA as an anti-inflammatory compound [4,5,19]. Nevertheless, this is the first *in vivo* analysis in mice that measures the CBDA effect on neuroinflammation. Even more, only one study before has aimed to examine the modulation of peripheral inflammatory responses by CBDA *in vivo* [19]. The pro-inflammatory profile of CBDA in PFC could be a factor contributing to its lack of effect on cocaine’s rewarding effects in the CPP paradigm. Anyway, the evaluation of other markers related to neuroinflammatory processes would help to better elucidate the possible CBDA modulation of immune response.

## CONCLUSIONS

Taken together, the results of the present study suggest that CBDA (0.001-1 mg/kg) has limited effects modulating mice behaviour. Only nociceptive threshold and despair-like behaviour were influenced by the administration of this phytocannabinoid. Surprisingly, CBDA exerted a pro-inflammatory profile in PFC. Discrepancies between the results presented in this study and previous literature highlights the need for more research in the study of this poorly understood phytocannabinoid.

## Abbreviations

5-HT_1A_: Serotonin receptor 1A
CBD: Cannabidiol
CBDA: Cannabiodiolic acid
COX-2: Ciclooxigenase-2
CPP: Conditioned place preference
EPM: Elevated plus maze
IL-6: Interleukin-6
PFC: Prefrontal cortex
PPAR: Proliferator-activated receptor
TRPA1: Transient receptor potential cation channel ankyrin 1
TRPM8: Transient receptor potential cation channel melastatin 8
TRPV1: Transient receptor potential cation channel vanilloid 1
TST: Tail suspension test

## ACKNOWLEDGEMENTS

This study was supported by the Ministerio de Economia y Competitividad (SAF2016-75966-R-FEDER and FPI grant to LAZ BES-2017-080066) and Ministerio de Sanidad (Retic-ISCIII, RD16/017/010), Plan Nacional sobre Drogas (2018/007). The Department of Experimental and Health Sciences (UPF) is a “ Unidad de Excelencia María de Maeztu” funded by the AEI (CEX2018-000792-M).

